# EEG of the dancing brain: Decoding sensory, motor and social processes during dyadic dance

**DOI:** 10.1101/2024.12.17.628913

**Authors:** Félix Bigand, Roberta Bianco, Sara F. Abalde, Trinh Nguyen, Giacomo Novembre

## Abstract

Real-world social cognition requires processing and adapting to multiple dynamic information streams. Interpreting neural activity in such ecological conditions remains a key challenge for neuroscience. This study leverages advancements in de-noising techniques and multivariate modeling to extract interpretable EEG signals from pairs of participants (male-male, female-female, and male-female) engaged in spontaneous dyadic dance. Using multivariate temporal response functions (mTRFs), we investigated how music acoustics, self-generated kinematics, other-generated kinematics, and social coordination uniquely contributed to EEG activity. Electromyogram recordings from ocular, face, and neck muscles were also modeled to control for artifacts. The mTRFs effectively disentangled neural signals associated with four processes: (I) auditory tracking of music, (II) control of self-generated movements, (III) visual monitoring of partner movements, and (IV) visual tracking of social coordination. We show that the first three neural signals are driven by event-related potentials: the P50-N100-P200 triggered by acoustic events, the central lateralized movement-related cortical potentials triggered by movement initiation, and the occipital N170 triggered by movement observation. Notably, the (previously unknown) neural marker of social coordination encodes the spatiotemporal alignment between dancers, surpassing the encoding of self-or partner-related kinematics taken alone. This marker emerges when partners can see each other, exhibits a topographical distribution over occipital areas, and is specifically driven by movement observation rather than initiation. Using data-driven kinematic decomposition, we further show that vertical bounce movements best drive observers’ EEG activity. These findings highlight the potential of real-world neuroimaging, combined with multivariate modeling, to uncover the mechanisms underlying complex yet natural social behaviors.

**Significance statement:** Real-world brain function involves integrating multiple information streams simultaneously. However, due to a shortfall of computational methods, laboratory-based neuroscience often examines neural processes in isolation. Using multivariate modeling of EEG data from pairs of participants freely dancing to music, we demonstrate that it is possible to tease apart physiologically established neural processes associated with music perception, motor control, and observation of a partner’s movement. Crucially, we identify a previously unknown neural marker of social coordination that encodes the spatiotemporal alignment between dancers, beyond self- or partner-related kinematics alone. These findings highlight the potential of computational neuroscience to uncover the biological mechanisms underlying real-world social and motor behaviors, advancing our understanding of how the brain supports dynamic and interactive activities.

## Introduction

A central challenge in neuroscience is understanding how the brain supports natural behavior in real-world contexts. Neuroimaging studies have traditionally been limited by bulky, motion-sensitive equipment, restricting research to controlled, motionless behaviors. This approach fails to capture how the brain manages the dynamic, multifaceted demands of everyday life, where cognition involves simultaneous neural processes, unconstrained movement, and interaction with ever-changing sensory environments—factors that traditional lab studies are poorly equipped to address (Stangl et al., 2023). Despite the recent advancements in mobile neuroimaging techniques (Niso et al., 2023) and algorithms for removing motion artifacts (Kothe and Jung, 2016), studying brain activity during natural behavior remains underexploited. As a result, it remains unclear how neural processes identified in lab-controlled studies generalize to real-world experiences, limiting our ability to interpret neural signals recorded during free behavior.

Here we used human collective dance as a model to study the neural basis of real-world interactions. We reason that dance offers an ideal testbed for several reasons: 1) it is culturally ubiquitous, hence broadly generalizable (Mithen, 2006; Dunbar, 2012); 2) it is complex yet controllable through musical structure (D’Ausilio et al., 2015); and 3) it encapsulates several intertwined neural processes, including auditory-tracking of music, movement control, monitoring others’ movements, and integrating these signals into cohesive experiences (Foster Vander Elst et al., 2023). These processes—notably targeting a variety of sensory and motor systems—can be effectively measured, for example, using electroencephalography (EEG). Yet, the main analytical challenge lies in disentangling these simultaneous neural signals (capturing sensory, motor, and social functions) from each other, and artifactual signals.

We tackled this challenge using multivariate temporal response functions (mTRFs), a computational approach that models the influence of different input variables on neural activity (Lalor et al., 2009; Crosse et al., 2016). We applied this method to a dataset of 80 participants, forming 40 dyads, who danced spontaneously to music while their brain activity, muscle activity, and full-body movements were recorded. Specifically, we captured EEG (64 channels), 3D full-body kinematics (22 markers), electrooculography (EOG), and electromyography (EMG, from neck and facial muscles), across various experimental conditions—detailed below (Bigand et al., 2024). mTRFs were meant to isolate four concurrent neural processes: 1) auditory perception of music, 2) motor control of specific body parts or specific movements, 3) visual perception of a partner’s body movements, and 4) visual tracking of social coordination, defined as the spatiotemporal alignment of movements between dancers, whether in-phase or anti-phase. Importantly, EOG and EMG signals were included as model predictors to account for potential muscle artifacts affecting the neural data. Additionally, we used event-related potential (ERP) analyses to anchor our findings in established physiological markers of sensory (auditory and visual evoked potentials) and motor (movement-related cortical potentials) processes (Novembre et al., 2018; Bach and Ullrich, 1997; Deecke et al., 1969).

Previous studies have used mTRFs to extract neural tracking of ecological auditory and visual stimuli, such as speech, music, or films (Di Liberto et al., 2015, 2020; O’Sullivan et al., 2017; Fiedler et al., 2019; Jessen et al., 2019; Bianco et al., 2024; Desai et al., 2024). However, aside from one human study and recent animal research incorporating body kinematics (Musall et al., 2019; Stringer et al., 2019; Di Liberto et al., 2021; Mao et al., 2021; Tremblay et al., 2023; Lanzarini et al., 2025), human studies that concurrently examine both sensory and motor processes using mTRFs—particularly in naturalistic behaviors—remain scarce. Furthermore, to our knowledge, no study has explicitly modeled social processes or addressed the neural activity associated with body-movement artifact leakage, as we do here. As such, our holistic approach aims to demonstrate that naturalistic human behaviors—implying real-time adaptation and movement—can be effectively explored using traditional electrophysiology. Therefore, our study highlights the potential of advanced neural analysis techniques to bridge the gap between lab-controlled and real-world neuroimaging research, enhancing our understanding of the neural basis of natural human behavior.

## Materials and methods

The EEG, EOG, EMG, and kinematic data analyzed here were collected as part of a previous study (Bigand et al., 2024), where participant dyads engaged in spontaneous dance under a 2×2 experimental design (Figs. 1a and 1b). The manipulated within-dyad factors were musical input (whether participants danced to the same [synchronous] or different [asynchronous] music) and visual contact (whether participants could see or not see their dancing partner).

**Figure 1.**
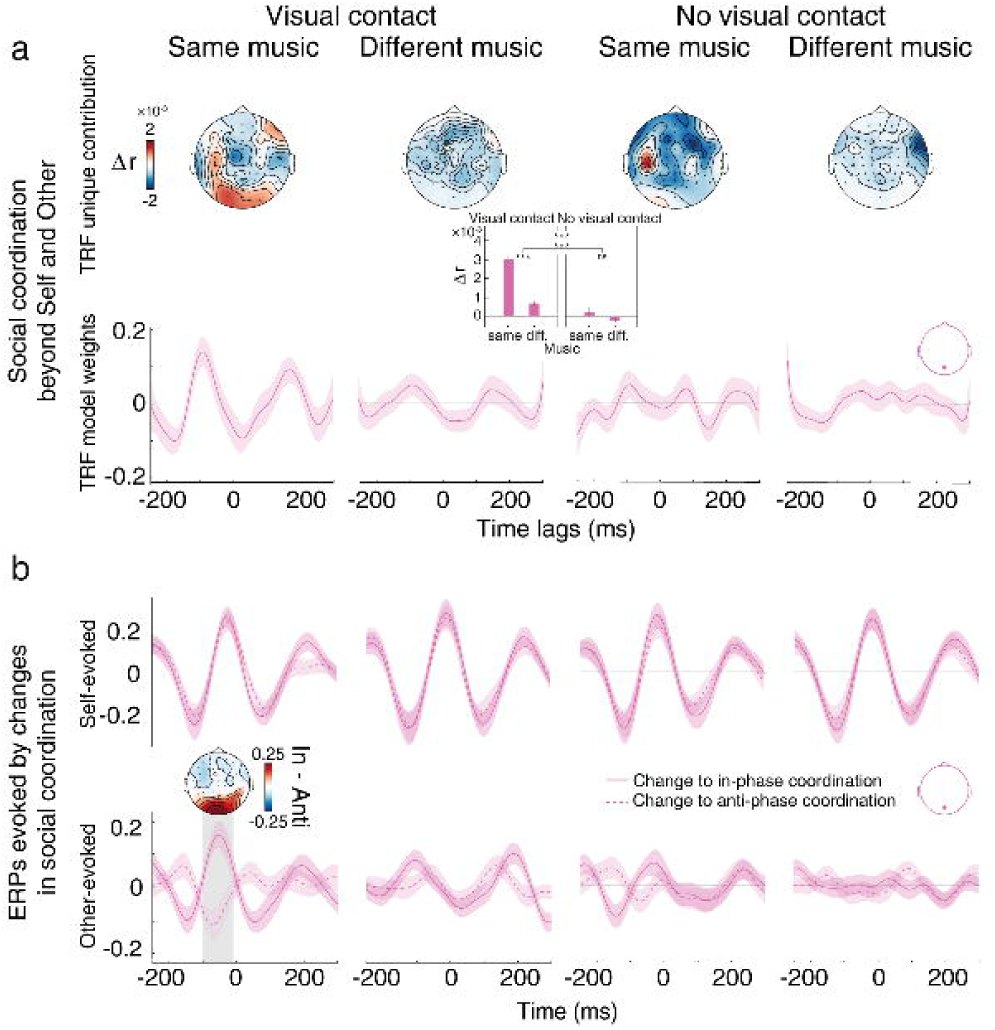
Experimental materials and methods. **a**, Experimental setup. We applied the mTRF method to a previously collected dataset (Bigand et al., 2024) for which dyads of participants danced spontaneously in response to music while we recorded electroencephalography (EEG, 64 channels), electrooculography (EOG), electromyography (EMG, from neck and face muscles), and 3D full-body kinematics (22 markers). **b,** Experimental design. Data were collected under the experimental conditions of the original study, which utilized a 2×2 factorial design. The two manipulated factors were musical input (whether participants listened to the same or different music presented through earphones) and visual contact (whether participants could see or not see each other). **c,** Overview of the modeling paradigm. We estimated multivariate Temporal Response Functions (mTRFs), which learned the optimal linear mapping between the set of variables of interest (here music, self- and other-generated movements, social coordination, as well as other control variables (not shown) such as ocular, facial and neck muscle activity) and the EEG data. **d,** Model comparisons. To assess the unique contribution of each variable (regressor) to the EEG data, we trained reduced models encompassing all variables apart from the specified one. The difference in prediction accuracy between the reduced and full model (encompassing all variables), denoted Δr, yields the unique contribution of this variable.

## Participants

80 participants (54 females; mean age: 26.15 years, SD: 6.43 years, 74 right-handed) formed 40 dyads (52% female-male, 41% female-female, and 7% male-male). To minimize inter-individual variability while maximizing generalizability, we recruited only laypersons (i.e. individuals without formal dance training). All participants forming a dyad were familiar with each other and were informed about the social nature of the task during recruitment, when they received the following message (translated from Italian): “You will have to come with someone you know (friend, family member, colleague…) with whom you will dance while listening to music (almost) like in a disco!”. As a measure of participants’ inclination toward social dance, we present the results of a post-hoc questionnaire. Specifically, participants rated the statement *“How often do you dance to music”* with a mean score of 4.363 (SD = 1.052) on a 6-point Likert scale (*1 = Never* to *6 = Very frequently*), and rated the statements *“When at a party, I am likely to be one of the first people dancing”* and *“I do not worry what other people think of my dancing skills”* with mean scores of 3.863 (SD = 1.626) and 3.863 (SD = 1.349), respectively, on a 6-point Likert scale (*1 = Strongly disagree* to *6 = Strongly agree*). Participants had normal or corrected-to-normal vision, normal hearing, and no history of neurological disorders. Data from five dyads were excluded due to recording failure in the motion capture system, leaving data from 70 participants for the analysis. All participants provided written informed consent to participate in the study and were compensated €25 for their participation. All experimental procedures were approved by “Comitato Etico Regionale della Liguria” (794/2021 - DB id 12093) and were carried out under the principles of the revised Helsinki Declaration.

## Musical stimuli

The musical stimuli consisted of eight songs with an average duration of 39.8 seconds (standard deviation: 1.95 seconds). Each song was presented in all four experimental conditions (see Experimental Design and Procedure below), resulting in a total of 32 trials. These songs were remakes of famous song refrains from electronic dance music and disco-funk genres (see Bigand et al. (2024)). Each song was adapted using the same four musical instruments: drums, bass, keyboards, and violin (the latter providing the vocal melody). All stimuli followed a 4/4 meter and spanned 20 bars. To create these adaptations, author FB and a professional composer (Raoul Tchoï) transcribed the original 4-bar refrain loops into MIDI format and synthesized them using MIDI instruments in Logic Pro X (Apple, Inc.). The rearranged songs were then systematically structured by repeating the 4-bar loops five times and sequentially adding each instrument to the musical scene, in the following order: (1) drums, (2) bass, (3) keyboards, (4) voice, (5) voice *bis* (i.e. the loop with full instruments was repeated twice). Loudness level across songs was controlled within a range of 1.5 LUFS (a measure accounting for the frequency sensitivity of the human auditory system). The songs were presented to the two participants forming each dyad through two separate EEG-compatible earphones (Etymotic ER3C), each connected to a distinct output channel of an audio interface (RME Fireface UC).

Every trial consisted of one song flanked with a fast-rising tone (rise time: 5 ms, fall time: 30 ms, frequency: 494 Hz, duration: 350 ms), preceded by 8 seconds of silence and followed by 7 seconds of silence, following this pattern: beep-silence-song-silence-beep. Trials were controlled using Presentation software (Neurobehavioral Systems), with synchronization between song presentation, EEG, and motion capture recordings achieved via TTL pulses. A TTL pulse was sent at the start of each trial from Presentation to both the EEG system (BioSemi ActiveTwo) and the motion capture system (Vicon; Lock+). This pulse activated the motion capture system, initiating recording, which automatically stopped after one minute. Simultaneously, the TTL pulse was stored alongside the continuous EEG recordings, which remained uninterrupted throughout the experiment. The TTL pulse, whose value varied based on the trial condition, enabled us to epoch the EEG data and retrieve the corresponding trial condition for analysis.

## Experimental design and procedure

EEG and kinematic data were recorded across four conditions derived from a 2×2 experimental design with visual contact (Yes, No) and musical input (Same, Different) as within-dyad factors. Conditions with or without visual contact were defined by the presence or absence of a curtain between the two participants in each dyad. Musical input was manipulated by presenting either identical or different songs to the participants through earphones. Because each song had a different tempo, the songs played simultaneously to the two participants were either perfectly synchronized (in the same-music condition) or slightly out of sync (in the different-music condition). To minimize inter-trial variability, this degree of asynchrony was maintained constant across trials belonging to the different-music condition (i.e. relative tempo difference between the two songs was precisely 8.5%). This was achieved by presenting participants with songs from different genres during the different-music condition (the tempo associated with electronic dance music songs was on average faster than that of disco-funk songs; see Bigand et al. (2024)). Trials were organized into four blocks, with each block including eight trials (two trials per condition, each trial featuring a different song). The presentation order of the blocks – and trials within blocks – was randomized, except for the deliberate presentation of subsequent pairs of yes-vision or no-vision trials to minimize the displacement of the curtain.

Before the experiment began, participants were told to behave as in a “silent disco,” in which they should face each other, enjoy the music, and remain still during periods of silence before and after the music. To enhance participants’ comfort, the overhead lighting was dimmed using alternating red and blue colored filters, creating a softer, “disco-like” atmosphere. Additionally, the experimenter remained out of sight in a custom-built cabin (enclosed by 1.5m-high panels), ensuring mutual invisibility between the experimenter and participants, as well as concealing the acquisition computers. Participants completed two training trials using songs not included in the main experiment to familiarize themselves with the task and setting. During this phase, they could request volume adjustments to their earphones, which were instructed to be set “as loud as possible without discomfort.” Participants were allowed (but not required) to dance freely within their designated space, keeping their head orientation towards their partner as steady as possible. Speaking or singing during trials was prohibited. Throughout the experiment, participants stood facing each other, each positioned within a marked area of 0.5×0.7 meters, with a separation of 2.5 meters between them.

## Kinematics data acquisition and preprocessing

3D full-body kinematics were recorded using wearable markers (22 per participant, size=14 mm). Markers were placed on specific body parts, denoted as follows (L = left, R = right, F = front, B = back): (1) LB Head, (2) LF Head, (3) RF Head, (4) RB Head, (5) Sternum, (6) L Shoulder, (7) R Shoulder, (8) L Elbow, (9) L Wrist, (10) L Hand, (11) R Elbow, (12) R Wrist, (13) R Hand, (14) Pelvis, (15) L Hip, (16) R Hip, (17) L Knee, (18) L Ankle, (19) L Foot, (20) R Knee, (21) R Ankle, (22) R Foot (Fig. 1a). Additionally, one supplementary marker was placed asymmetrically on either the left or right thigh of each participant. This marker was only used to facilitate Nexus software in the distinction between participants and was not considered in subsequent analyses. Eight optical motion capture cameras (Vicon system) recorded the markers’ trajectories at a sampling rate of 250 Hz. The cameras were positioned to capture the participants from various angles, ensuring that each participant was visible to at least six cameras even when visual contact was obstructed by the curtain. A high-definition video camera, synchronized with all the optical motion capture cameras, recorded the scene from an aerial view (Vicon Vue; 25 Hz sampling frequency; 1,920 × 1,080 pixels). We used a Vicon motion capture system to record full-body 3D positions with high spatial (<1 mm precision) and temporal (250 Hz) resolution. While alternative methods, such as inertial measurement units or accelerometers, could be considered, the feasibility of repeating our study with fewer markers remains to be tested. Notably, full-body tracking was essential here for breaking down complex dance kinematics into the elementary movement components that drove neural signals (see Kinematic feature selection below).

Markers’ trajectories were corrected for swaps or mislabels via the Nexus manual labeling tool (Vicon). Then, automated correction of frequent and systematic marker swaps was performed using custom Python code. Any gaps in the marker trajectories were then filled using the automatic gap-filling pipeline in Nexus. The proportion of time with gaps, calculated for each marker and averaged across participants, ranged from a minimum of 0.128% (L Foot) to a maximum of 2.767% (R Hip), with a mean of 0.688% and a standard deviation of 0.739%. Lastly, all trajectories were inspected visually within Nexus software and manually adjusted if they did not match the aerial-view video recording. Subsequent data analyses were carried out in Python using custom code. Marker trajectories comprised 3D positions (along x, y, and z axes) corresponding to each of the 22 body parts, resulting in time-series of posture vectors of 66 dimensions.

## EEG data acquisition and preprocessing

We recorded neural activity from both participants simultaneously using a dual-EEG setup with the BioSemi ActiveTwo system. This setup consists of two AD-Boxes, each independently recording and referencing EEG from a single participant. The data from the two AD-Boxes are synchronized at the hardware level: the ‘slave’ AD-Box transmits data via optical fiber to the ‘master’ AD-Box, which then relays all EEG signals and triggers information to the acquisition computer. For a detailed schematic of the BioSemi ActiveTwo dual-EEG configuration, see Barraza et al. (2019). For each participant, the EEG was recorded from 64 Ag/AgCl active electrodes (placed on the scalp according to the extended international 10–10 system). To help retain the naturalistic nature of the study, we used 2-meter-long cables, custom-built by the manufacturer to meet our specific requirements. Each EEG amplifier was positioned behind the participant at hip height, with cables taped to the upper back to minimize weight while ensuring they remained loose enough to prevent any perceived constraint or pulling. This setup allowed participants to move relatively freely while remaining within their designated area (see Experimental Design and Procedure).

EEG signals were digitized at 1024 Hz using the BioSemi Active Two system. Subsequently, the data were pre-processed and analyzed using Matlab R2022. Measuring EEG from moving participants is susceptible to muscular artifacts in the recordings. To mitigate this issue, we pre-processed the EEG data of the dancing participants using a fully data-driven pipeline that we had previously developed for analyzing EEG data in awake monkeys (Bianco et al., 2024). This pipeline primarily utilizes open-source algorithms from Fieldtrip (Oostenveld et al., 2011) and EEGLAB (Delorme and Makeig, 2004) toolboxes. EEG signals were digitally filtered between 1 and 8 Hz (Butterworth filters, order 3), down-sampled to 100 Hz, and trimmed according to the duration of the trial-specific songs. Faulty or noisy electrodes were provisionally discarded before re-referencing the data using a common average reference. This was done to prevent the leakage of noise to all electrodes during re-referencing. Criteria for flagging faulty or noisy electrodes included prolonged flat lines (lasting more than 5 seconds), abnormal inter-channel correlation (lower than 0.8), or deviations in amplitude metrics from the scalp average (mean, STD, or peak-to-peak values exceeding 3 STD from the scalp average). These assessments were made using EEGlab’s *clean_flatlines* and *clean_channels* functions (Delorme and Makeig, 2004) and custom Matlab code. To remove movement artifacts, we further denoised the re-referenced data using a validated algorithm for automatic artifact correction: Artifact Subspace Reconstruction (ASR, threshold value 5) (Kothe and Jung, 2016). This algorithm has been previously applied to human data, including in music-making and dance studies (Ramírez-Moreno et al., 2023; Theofanopoulou et al., 2024). Finally, eye-movement artifacts were subtracted from the ASR-cleaned data using another automatic artifact-correction algorithm – ICA, using EEGlab’s *IClabel* function (Pion-Tonachini et al., 2019). Independent Components that were classified by *IClabel* as eye-movement artifacts (i.e., those for which the ‘eye’ category had the highest probability, with no minimum threshold) were removed. At this stage, noisy or faulty electrodes (as assessed at the start of this preprocessing pipeline) were interpolated by replacing their voltage with the average voltage of the neighboring electrodes (20-mm distance).

## EOG and EMG data acquisition and preprocessing

Two EOG channels were recorded using surface Ag–AgCl electrodes from all participants. Electrodes were attached using disposable adhesive disks at specific anatomical locations: the left and right outer canthi. Additionally, we also recorded four EMG signals from the cheeks (the left and right zygomata) and the neck (the left and right paraspinal muscles) for control purposes. EOG/EMG signals were digitized at 1024 Hz using the BioSemi Active Two system. The EOG/EMG data were filtered, down-sampled, and trimmed similarly as the EEG data, re-referenced using scalp average, and ASR-cleaned using a threshold value of 5, to maintain consistency with the EEG signals from scalp channels. It should be noted that these signals were measured from a subset of participants (n=58), therefore all subsequent analyses involving this data subset include only these participants.

## Multivariate temporal response functions (mTRFs)

Events, such as hearing a fast-rising sound or initiating a movement, elicit phase-locked brain activity within a specific time window [t1,t2], which can include post-event (e.g., response to sounds) and pre-event (e.g., movement initiation) components (Luck, 2014). Temporal response functions (TRFs) can be used to characterize this relationship at the level of EEG electrodes (Lalor et al., 2009; Crosse et al., 2016). In this study, we applied mTRFs to delineate the distinct neural processes that occur simultaneously during dyadic dance. Specifically, we first extracted a diverse set of time-resolved variables, representing: (I) musical input, (II) self-generated movements, (III) partner-generated movements, (IV) social coordination, and (V, VI, and VII) ocular, facial and neck muscle activity (Fig. 1c, left). Next, we estimated TRFs (Fig. 1c, middle) to quantify how these variables modulate EEG signals, for each electrode separately (Fig. 1c, right). The following sections provide a detailed explanation of these two steps.

*Step 1: Extraction of variables*. *(I) Music*. Musical input was represented using spectral flux, which captures fluctuations in the acoustic power spectrum. Spectral flux has been shown to outperform other acoustic features, such as the envelope and its derivative, in predicting neural signals elicited by music (Weineck et al., 2022). To extract it, we first bandpass filtered the musical stimuli into 128 logarithmically spaced frequency bands ranging from 100 to 8000 Hz using a gammatone filter bank. Spectral flux was then computed for each frequency band by calculating the first derivative of the band’s amplitude over time. Finally, the broadband spectral flux, representing overall changes in the spectral content, was derived by averaging the spectral flux across all 128 bands. *(II and III) Self- and other-generated movements*. The movements produced by each participant (self-generated) and their partners (other-generated) were represented using velocity magnitude. To reduce dimensionality, full-body trajectories were decomposed into 15 principal movement patterns that collectively explained over 95% of the kinematic variance (see Methods in Kinematic feature selection below). The velocity magnitude of each principal movement was calculated by taking the first derivative of its position over time and then computing the absolute value of this derivative. Out of the 15 principal movements, preliminary analyses identified bounce as the movement that explained most of the neural encoding of both self-and other-generated movements (see results in Kinematic feature selection below, and Fig. 2). Consequently, only the velocity magnitude of the bounce trajectory was included in subsequent models. *(IV) Social coordination*. To assess social coordination, we extracted a categorical measure to determine whether the bounce movements of participants within a dyad were in-phase or anti-phase. This measure indexed whether both individuals moved in the same direction (in-phase) or opposite directions (anti-phase). We obtained this measure by multiplying the signs of the bounce velocity time series (i.e., the respective directions of movement) across the two participants forming a dyad. *(V, VI, and VII) Ocular, facial, and neck muscle activity.* To control for muscular activity potentially leaking into the EEG signals, we also included EOG (measured from the left and right eyes) and EMG (measured from the cheeks and the neck) time-series in the models. Both EOG channels (left and right eye) were included to capture horizontal saccades, which generate opposite left-right activity (positive values on one side and negative on the other). For cheek and neck muscles, the average signal from the left and right EMG channels was used, respectively. All these variables, each of which is time-resolved, were down-sampled to match the EEG sampling frequency of 100 Hz and trimmed according to the duration of the trial-specific songs. To account for inter-individual variability, all variables were standardized on a per-participant basis by normalizing each time-series to its standard deviation across all trials for the corresponding participant.

**Figure 2.**
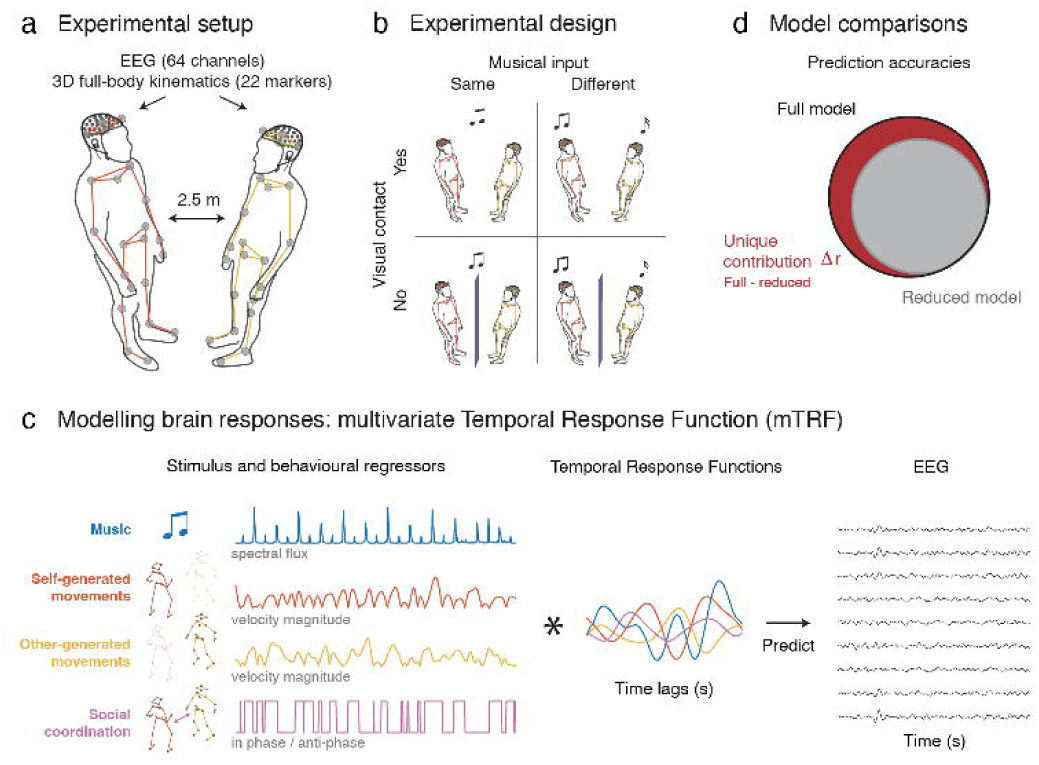
Neural encoding of self- and other-related movements across different principal movements (PMs). Bars represent the unique contribution (Δr) of each PM (grand-average) to the EEG signal recorded from the self (electrode Cz) or the other (electrode Oz). Δr values represent the difference in EEG prediction accuracy between the PM-specific reduced models and the full model, for self- and other-generated movements, respectively. Error bars represent ±1 standard error mean (SEM). Gray circle diagrams illustrate the proportion (%) of kinematic variance explained by each PM, with the first 15 PMs accounting for more than 95% of the total variance. Together, these results indicate that bounce (PM10) was the strongest predictor of EEG activity, whether self-generated or observed in others, despite accounting for only ∼1% of kinematic variance.

*Step 2: mTRF estimation*. We estimated TRFs via a multivariate lagged regression, which fitted the optimal linear mapping between the abovementioned variables and EEG at each electrode (mTRF toolbox, encoding model (Crosse et al., 2016); Fig. 1c). A time-lag window of -250 to 300 ms was selected to encompass commonly observed ERP responses associated with sound perception (Novembre et al., 2018), execution of fast-repeated movements (Gerloff et al., 1997), and visual perception of biological movements (Jokisch et al., 2005). This window also ensured that the contribution of redundant (potentially irrelevant) information was minimized, especially considering the rhythmic structure of the task, with musical beats and some dance movements (e.g., bounce) occurring approximately every 500 ms. Importantly, in a control analysis we confirmed that the selected window did not reduce prediction accuracy compared to a broader [-700, +700 ms] window. For each participant and experimental condition, mTRFs were estimated, including either all variables simultaneously (full model) or all variables except the specified one (reduced models; see below for details). Participant- and condition-specific TRFs were estimated as the average TRF required to predict each of the eight condition trials using data from the remaining seven trials (i.e., TRFs were fit eight times). Regularized (ridge) regression was used to fit the TRFs, maximizing prediction accuracy (the correlation between the predicted and actual EEG data; Pearson’s *r*) without overfitting the training data. The optimal regularization parameter (λ) was selected via leave-one-out cross-validation across trials (i.e., songs), tested over a range from 0 to 10D (0, 10DD, 10D³, …, 10D). This yielded one optimal λ value per trial, condition, and participant. Finally, prediction accuracies for each condition were assessed using a generic approach (Di Liberto and Lalor, 2017; Jessen et al., 2019), where the Pearson’s *r* between predicted and actual EEG data was calculated across all eight trials of the *n*^th^ participant, with predictions based on a generic TRF averaged across the subject-specific TRFs of the *N-1* remaining participants (N=70). The prediction accuracy of a model describes the amount of EEG variance that the model can account for. To evaluate the unique amount of EEG variance that each variable accounts for, we constructed reduced models that included all variables apart from the specified one. The difference in prediction accuracy between the full (comprising all variables) and the reduced model yielded the unique contribution, denoted as Δ*r*, of that specific variable to the variance explained in the EEG data (Fig. 1d).

## Kinematic feature selection

To reduce dimensionality, we used a data-driven method to determine a subset of kinematic variables to use in the TRF models. The kinematic data were decomposed into a set of principal movements using Principal Component Analysis (PCA), following the same pipeline as described in Bigand et al., (2024). These principal movements reflect movement primitives that are generalizable across trials, conditions, and participants. This PCA approach has been previously validated for a wide range of human movements, including dance (Troje, 2002; Daffertshofer et al., 2004; Toiviainen et al., 2010; Federolf et al., 2014; Yan et al., 2020; Bigand et al., 2021). The first 15 principal movements – accounting for more than 95% of the kinematic variance – were retained for further analyses (Federolf et al., 2014; Bigand et al., 2021). The score time series obtained from the PCA reflected the position of each principal movement over time. These time-series were low-pass filtered below 6 Hz using a Butterworth filter (second-order, zero-phase) to increase the signal-to-noise ratio. These 15 principal movements were reminiscent of common “dance moves” such as body sway, twist, upper-body side bend and rock, bounce, side displacement, head bob, hip swing, and hand movements (see Fig. 2 and Video 1) (Bigand et al., 2024).

To determine which principal movements to include in the TRF models, we tested their association with EEG modulations. Previous evidence suggests that TRFs or equivalent models can accurately capture neural activity associated with both the generation (Musall et al., 2019) and the observation (O’Sullivan et al., 2017; Jessen et al., 2019) of biological movement. Accordingly, we tested the unique contribution of the 15 principal movements, either self-generated or generated by (and therefore observed in) the dancing partner. In other words, we fit 30 reduced models and computed the difference in prediction accuracy (Δ*r*) between each reduced model and a full model (including all 30 principal movements plus spectral flux) for each participant and condition, using the generic approach outlined above (see Methods in mTRF estimation above). Spectral flux was included in the full model to ensure that the explanatory power of individual principal movements was not influenced by movements correlated with the music as participants were dancing to music. To reduce computational cost, the 15 models for other-generated movements were trained in the visual conditions, while those for self-generated movements were trained in the non-visual conditions. This ensured a balanced number of trials for analyzing both motor control and movement observation activities while testing movement observation under the conditions where it was most likely to occur. The full model was trained across all conditions, allowing for the computation of “self” and “other” Δ*r* values, averaged across the two non-visual and visual conditions, respectively.

The results of this preliminary analysis revealed that bounce movement—i.e., vertical oscillations of the body achieved through knee flexion and extension—was largely the main contributor to EEG prediction, notably across both self- and other-generated movements (see PM10; Fig. 2), despite accounting for no more than 1% of the total kinematic variance. Specifically, self-generated bounce alone explained >84% of the EEG prediction gain (Δ*r* > 0) across all principal movements at electrode Cz, commonly associated with motor activity (Kornhuber and Deecke, 1965; Deecke et al., 1969; Shibasaki et al., 1980; Smulders and Miller, 2011; Vercillo et al., 2018). Additionally, other-generated bounce alone accounted for >80% of EEG prediction gain at electrode Oz, a canonical site indicative of motion-evoked visual responses (Kubová et al., 1995; Bach and Ullrich, 1997; Puce et al., 2000; Jokisch et al., 2005; O’Sullivan et al., 2017). Given these results, bounce will serve as the primary movement feature in all subsequent analyses. Henceforth, when referring to “movement” in the following sections, we specifically denote “bounce” (except when discussing the results of the body-part-specific analyses).

## Statistical analyses

We assessed the distinct neural encoding of music, self-generated movements, other-generated movements, and social coordination while controlling for artifact leakage from eye, facial, and neck movements. We created seven reduced models, accounting for: (I) music (all variables minus spectral flux); (II) self-generated movements (all variables minus velocity magnitude of self-generated bounce); (III) other-generated movements (all variables minus velocity magnitude of other-generated bounce); (IV) social coordination (all variables minus interpersonal bounce coordination); and (V, VI and VII) ocular, facial and neck muscle activity (all variables minus EOG, facial EMG, or neck EMG, respectively). We then compared the prediction accuracies of these reduced models to that of a full model encompassing all seven variables, i.e., the unique contribution Δ*r* of each variable.

To compare the unique contributions of music, self-generated movements, other-generated movements, and social coordination across different experimental conditions (visual contact (yes/no) x music (same/different)), Δ*r* values were averaged across relevant electrodes for each participant and predictor. Relevant electrodes were defined independently for each predictor as those that exhibited a prediction gain (Δ*r* > 0). For each predictor, this gain was computed across conditions where the associated neural process was expected to occur: all conditions for music and self-related movements, and visual conditions for other-generated movements and coordination. The Δ*r* values to be statistically compared were computed for each condition and then averaged across the defined electrodes. This yielded a Δ*r* value per participant, condition, and variable (music, self- and other-generated movements, and social coordination).

We assessed differences in unique contributions across conditions using a 2×2 repeated-measures ANOVA with the factors “visual contact” and “musical input”. Δ*r* values were normally distributed and entered into the ANOVA as the dependent variable. To control for multiple comparisons across the four variables, p-values were Bonferroni-corrected.

## Event-related potentials (ERPs)

### Extraction of ERPs

To aid in interpreting the TRF results—particularly the physiological origins of the TRF model weights—we examined phase-locked neural responses, i.e., event-related potentials (ERPs), evoked by changes in music intensity, self-generated movement velocity, other-generated movement velocity, and social coordination (transitions between in-phase and anti-phase coordination – see below). EEG responses are largely evoked by fast changes in the environment (Somervail et al., 2021), including fluctuations in the auditory spectrum (Weineck et al., 2022) and peaks in movement velocity (Varlet et al., 2023). Therefore, we determined the onset times of events, such as sounds or movements, by identifying peaks in the respective time series using Matlab’s *findpeaks* function (with default parameters). Acoustic onsets were thus aligned with musical notes played by any of the four instruments in the stimuli, while motion onsets were aligned with velocity peaks. Coordination onsets were obtained from the first derivative of the coordination time series, corresponding to transitions between in-phase and anti-phase states. To improve the signal-to-noise ratio, acoustic peaks were filtered by selecting only the most salient, i.e., those that were 3 STD away from the mean of the trial. This step was unnecessary in the case of the other variables, such as movement and coordination, presumably because the kinematic data had already been low-pass filtered, as described earlier. Consequently, the signal-to-noise ratio for these variables was already maximized. The ERP epochs spanned the same time window as the TRFs (−250 to 300 ms).

### ERP sensitivity to variables’ intensity

ERP amplitude largely depends on the differential intensity of the evoking change, and this sensitivity to differential intensity is supramodal, i.e., it’s a property observed across different sensory systems (Somervail et al., 2021). Here, to quantify whether ERPs were modulated by the differential amplitude of musical sounds or by the speed of self- and other-generated movements, we categorized acoustic onsets into *soft/loud* and movement onsets into *slow/fast*. For each participant and experimental condition, we selected acoustic and motion onsets with the highest and lowest 20% values of spectral flux or velocity magnitude, respectively. Similarly, to quantify the ERP modulation as a function of coordination, we grouped coordination onsets into their two possible values: change to *in-phase* or *anti-phase*. Following established ERP literature (Jokisch et al., 2005; Novembre et al., 2018; Vercillo et al., 2018), epochs linked to external stimuli (music and other) were baseline corrected using a pre-stimulus interval (−250 to 0 ms), while epochs involving internally-initiated actions (self and coordination) were baseline corrected using the entire epoch duration. Differences between the two groups (*soft* vs. *loud*, *slow* vs. *fast,* or *in-phase* vs. *anti-phase*) were tested separately for each experimental condition, by means of a cluster-based permutation test (implemented in Fieldtrip, with 1000 permutations [Maris and Oostenveld, 2007]). This analysis focused on the EEG channels of interest informed by the mTRF results: Fz (music), C3 and C4 (self-generated movements), Oz (other-generated movements), and Oz (social coordination).

## Body-part-specific mTRF (self)

In the main analysis, motor activity was assessed using mTRFs predicted by the kinematics of self-generated bounce, as this movement explained most motor activity across the 15 principal movements identified via PCA. Hence, the main analysis does not differentiate between body parts, as the bounce movement activates nearly all of them (see Fig. 2 and Video 1), making it challenging to determine whether the movement of specific body parts drove specific motor activities. To address this issue, we leveraged kinematic data from all parts of the body to calculate the unique contribution of self-generated motion to the EEG from the left and right hand, left and right foot, and head velocity magnitudes. Specifically, we created a full model that included major body markers (‘LB Head’, ‘LF Head’, ‘RF Head’, ‘RB Head’, ‘Sternum’, ‘L Shoulder’, ‘R Shoulder’, ‘L Hand’, ‘R Hand’, ‘Pelvis’, ‘L Hip’, ‘R Hip’, ‘L Knee’, ‘L Foot’, ‘R Knee’, ‘R Foot’) along with neck EMG controls, and five reduced models, each excluding specific markers: left/right hand markers, left/right foot markers, and the average of the four head markers. Markers expected to be almost intrinsically correlated with hand and foot movements (e.g., elbows, wrists, and ankles) were not included in the full model. As in the main analysis, the unique contribution of each body part’s kinematics to the EEG variance was determined by the difference in prediction accuracy between the full model and each reduced model.

## Encoding of social coordination

### Coordination beyond self and other?

Social coordination was operationalized as the spatiotemporal alignment of movements produced by participants (self-generated) and their partners (other-generated). Specifically, this construct assessed whether participants and their partners not only bounced *at the same time* but also *in the same direction*. As such, social coordination relied on both temporal features (velocity magnitude time-series) and spatial features (velocity sign time-series), with the latter indicating the up versus down phases of movement. In contrast, the measures of self and other were derived solely from temporal features. Consequently, it was essential to conduct a control analysis to assess the extent to which social coordination was influenced by the spatial characteristics of both self- and other-generated movements. To address this, we implemented an mTRF analysis utilizing a comprehensive model that incorporated music, self-generated and other-generated movements (velocity magnitude time-series), social coordination, and the spatial directions of both self- and other-generated movements (velocity sign time-series). For this control analysis, we did not include other control variables, such as muscular activity, because the previous analyses already demonstrated that these do not predict social coordination.

### Coordination ERPs driven by self or other?

In our main ERP analysis, we extracted “coordination ERPs” by epoching EEG time-series at transition onsets between in-phase and anti-phase coordination (see Extraction of ERPs methods described above). These transitions could potentially arise from changes in movement direction elicited by either the self or the partner. To disentangle these two possibilities—specifically, whether ERPs related to social coordination were driven by self-generated movements (self) or by partner-generated movements (other)—we categorized coordination ERPs into two distinct groups: those triggered by self-movements (i.e., when coordination changes were aligned with shifts in the velocity sign of self-generated movements) and those triggered by partner movements (i.e., when coordination changes aligned with shifts in the velocity sign of other-generated movements). We quantified the *in-phase/anti-phase* ERP modulation separately for these two groups, following the methods outlined previously (see previous ERP analyses). Differences between in-phase and anti-phase onsets were assessed independently for the “self” and “other” groups in each experimental condition using a cluster-based permutation test (implemented in FieldTrip with 1000 permutations [Maris and Oostenveld, 2007]). This analysis focused on the channel of interest informed by the mTRF results: Oz.

## Results

### Multivariate temporal response functions (mTRFs)

In our analysis, we assessed the unique contributions of four variables to the EEG by comparing the prediction gain (Δ*r*) between a full mTRF model and reduced models excluding each variable of interest (see Methods for details). The results showed that musical sounds, self-generated movements, other-generated movements, and social coordination each made distinct contributions to participants’ neural activity. This allowed us to isolate four neural processes co-occurring during dyadic dance: (I) auditory perception of music, (II) control of movement, (III) visual perception of the partner’s body movements, and (IV) visual tracking of social coordination. These processes were clearly distinguished from ocular, facial, and neck muscle artifacts (V, VI, and VII). The following section provides detailed information on each of these EEG activities.

***(I) Auditory perception of music.*** The spectral flux of the music uniquely predicted EEG across frontal and parietal electrodes, as evidenced by the prediction gain Δ*r* (the difference between the prediction of the full model and that of the reduced model excluding spectral flux) at each electrode (Fig. 3a). A repeated-measures ANOVA, with “musical input” and “visual contact” as factors, yielded a main effect of vision, demonstrating a significant reduction in the prediction gain Δ*r* when participants could see each other (*F*(1,57) = 7.48, *p* = .033; Fig. 4). This finding suggests a diminished neural tracking of music when participants could see their partners. The regression weights associated with the music TRF model (representing electrode Fz) highlight three post-stimulus modulations, i.e., a positive-negative-positive pattern with peaks at around +60, +120, and +200 ms post-stimulus, respectively (Fig. 3b). The weights also exhibit a consistent peak around -200 ms pre-stimulus, which, considering the periodic rhythmic nature of the music, is likely evoked by the preceding beat sound. We confirmed so by observing that the sound differential intensity (specifically, the spectral flux value) of the previous beat modulated the amplitude of this -200 ms peak.
***(II) Control of movement (self-generated).*** Self-generated movements uniquely predicted EEG across central and occipital electrodes, as indicated by the electrode-specific prediction gain Δ*r* (Fig. 3a). The ANOVA did not yield evidence of significant effects of musical input or visual contact on the unique contribution of self-generated movements on EEG signals, suggesting comparable motor control processes across conditions (all *ps* > .224; Fig. 4). The TRF weights associated with self-generated movements (representing the average between electrodes C3 and C4) highlighted three main modulations, i.e., a negative-positive-negative pattern with peaks at around -100, 0, and +80 ms relatively to movement onset, respectively (Fig. 3b).
***(III) Visual perception of partner’s body movements (other-generated).*** Other-generated movements uniquely predicted EEG across occipital electrodes, surrounding the visual cortex (Fig. 3a), only when participants could see each other. This was confirmed by the ANOVA, yielding a main effect of visual contact (*F*(1,57) = 83.23, *p* < .001; Fig. 4). This finding is consistent with the expectation that neural tracking of others’ movements can only occur when these movements are observable. The TRF weights associated with other-generated movements (representing electrode Oz) highlighted a biphasic modulation characterized by a positive peak at around +70 ms, and a negative peak at around +160 ms relative to movement onset (Fig. 3b).
***(IV) Social coordination.*** Social coordination uniquely predicted EEG primarily across occipital electrodes (Fig. 3a), especially when participants could see each other and listened to the same music. This was supported by the ANOVA, which yielded main effects of visual contact (*F*(1,57) = 249.75, *p* < .001) and musical input (*F*(1,57) = 30.22, *p* < .001), along with a significant interaction effect (*F*(1,57) = 50.10, *p* < .001) (Fig. 4). Follow-up comparisons revealed that EEG prediction accuracy was specifically enhanced when participants danced to the same music, but only with visual contact (Δ = 0.0009, *SE* = 0.0001, *p* < .001); no music effect was observed without visual contact (*p* = .676) (Fig. 4). This suggests that the level of coordination between participants is encoded in each participant’s EEG, and that neural tracking occurs primarily when partners are visible and synchronizing to the same musical tempo. In this condition, the TRF weights (representing electrode Oz) exhibited a quadriphasic pattern characterized by negative-positive-negative-positive peaks, at -180, -90, +30, and +160 ms relative to a change in coordination (between in-phase and anti-phase – see Methods), respectively (Fig. 3b).

**Figure 3.**
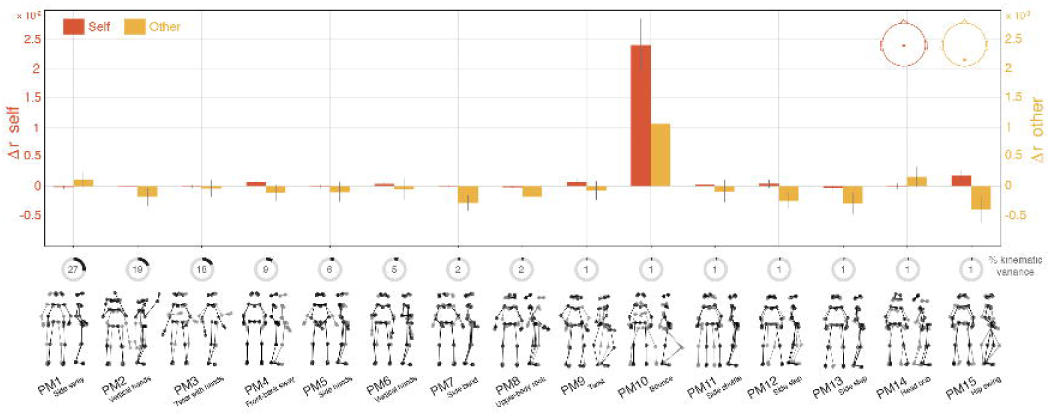
Distinct EEG activities related to music, self- and other-generated movements, social coordination, and muscle artifacts. **a,** Topographical maps of the unique contribution of each model variable to the predicted EEG. Δr topographical maps represent the grand-average difference in EEG prediction accuracy between the reduced models (excluding the variable of interest) and the full model (including all four variables, plus ocular, facial, and neck muscle activity; see Fig. 1d), for each EEG electrode and experimental condition. **b,** Ridge regression weights for TRFs corresponding to music (Fz), self-generated movements (averaged across C3, C4), other-generated movements (Oz), social coordination (Oz), and ocular (F8), facial (T8), and neck (Oz) muscle activity for the full-model TRF. Grand-average weights are shown. Shaded areas represent ±1 SEM.

**Figure 4.**
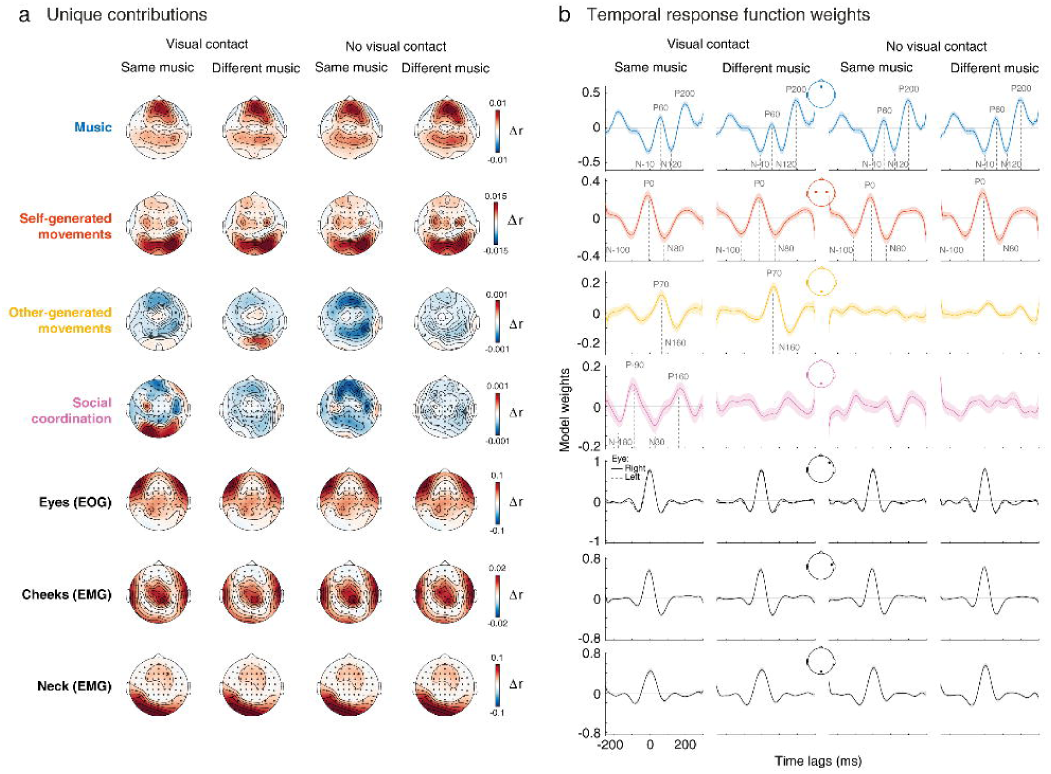
Comparison of unique contributions across experimental conditions. Bars indicate the grand-average unique contributions (averaged over electrodes showing a gain; Δr >0) of each model variable, across conditions. Error bars represent ±1 SEM. Stars indicate significant main effects of visual contact and musical input, as well as the interaction between the two factors (2×2 repeated measures ANOVA, Bonferroni-corrected; *p_bonf_<.05, **p_bonf_<.01, ***p_bonf_<.001).

***(V, VI, and VII) Ocular, facial, and neck muscle artifacts.*** EOG and EMG signals uniquely predicted EEG at electrode sites that closely matched artifactual topographical maps documented in previous EEG research (Fig. 3a) (Goncharova et al., 2003; Plöchl et al., 2012). Specifically, EMG from facial and neck muscles predicted EEG activity at the scalp periphery, which is typical of muscle contraction topographies (Goncharova et al., 2003), while EOG predicted EEG activity nearby the eyes (electrodes AF7 and AF8, approaching the lateral canthi), characteristic of eye saccades (Plöchl et al., 2012). Note that most blinks-related artifacts were presumably removed beforehand via ASR and ICA pipelines (see Methods). The TRF weights for eye, facial, and neck movements displayed features of instantaneous impulse responses (Fig. 3b), indicating that non-cerebral signals propagate to the EEG without measurable delay—a characteristic previously established for artifact leakage (Croft and Barry, 2000). Additionally, these EOG and EMG signals contributed orders of magnitude more to the EEG than brain processes (compare Δ*r* scales within Fig. 3a), another expected property of EMG and EOG activations (Urigüen and Garcia-Zapirain, 2015). Collectively, these findings underscore the efficiency of our analysis in distinguishing simultaneous neurophysiological processes from each other, as well as from movement-related artifact leakage.

### Event-related potentials (ERPs)

To elucidate the physiological origins of the temporal responses modeled by the mTRFs, we extracted ERPs by epoching the EEG time series around salient changes (see Methods) in music, self-generated movements, other-generated movements, and social coordination. This analysis was specifically designed to clarify the neurophysiological origin of the temporal responses, or model weights, modeled by the mTRFs. Notably, the focus was not on the condition-specific contribution of these responses to the EEG, as ERP analysis cannot fully account for concurrent contributions from other variables. Rather, the results demonstrated that the ERPs exhibited morphologies closely resembling the TRF weights observed earlier (compare Fig. 5 with Fig. 3b) and consistent with typical EEG markers of sensory and motor processes established in laboratory-controlled studies. Detailed ERP results for each process are presented in the following sections.

***(I) Auditory perception of music.*** The individual sounds embedded within the musical tracks elicited a characteristic triphasic ERP response, consisting of an early positivity (P50), followed by a widespread negativity (N100), and a later positivity (P200), all displaying a frontal topographic distribution (Fig. 5). This pattern aligns with established findings from both ERP (Novembre et al., 2018; Di Liberto et al., 2020) and TRF studies (Di Liberto et al., 2015, 2020; Fiedler et al., 2019; Jessen et al., 2019; Kern et al., 2022) in motionless participants, and closely resembles the regression weights of the music TRF observed in our study with dancing participants (Fig. 3b). These similarities suggest that our music TRF primarily captured phase-locked responses evoked by the individual sounds embedded within the musical stimuli, as observed in previous work (Di Liberto et al., 2020; Bianco et al., 2024). To validate this assumption, we further report a known physiological property of these responses—ERP amplitude sensitivity to variations in stimulus intensity (Somervail et al., 2021)—as evidenced by the amplitude of the P200 being larger in response to loud *vs* soft sounds (Fig. 5).
***(II) Control of movement (self-generated).*** ERPs time-locked to self-generated movements displayed a triphasic pattern, characterized by a pre-motor negativity (N-100), a positivity at movement onset (P0) and a post-motor negativity (N100), with a central distribution (Fig. 5). These components are reminiscent of movement-related cortical potentials (Shibasaki et al., 1980; Hallett, 1994), which might include steady-state movement-evoked potentials (Gerloff et al., 1997) or readiness potentials (Kornhuber and Deecke, 1965; Vercillo et al., 2018) (see also body-part-specific analyses, and Fig. 6). The pattern closely mirrors the regression weights observed in our TRF model of self-generated movements (Fig. 3b). Notably, the amplitude of the ERPs associated with self-generated movements was larger during relatively faster, as opposed to relatively slower movements, a pattern previously suggested to reflect increased motor activity during higher-rate movement execution (Brunia et al., 2011). These results suggest that the mTRF model effectively captured EEG potentials traditionally linked to motor control, with amplitudes modulated by movement speed.
***(III) Visual perception of partner’s body movements (other-generated).*** When the participants could see each other, the observed partner-generated movements elicited biphasic responses in occipital regions, characterized by a positive peak at ∼70 ms (P70) and a negative peak at ∼160 ms (N160) (Fig. 5). This pattern resembles traditional visual responses to biological motion, notably characterized by the N170 component, typically observed around 170 ms post-movement onset (Kubová et al., 1995; Bach and Ullrich, 1997; Puce et al., 2000; Jokisch et al., 2005). Similar to the responses associated with music and self-generated movements, these ERPs closely align with the regression weights yielded by the TRF model of other-generated movements (Fig. 3b). As for ERPs evoked by motor control (previous section), ERP amplitudes scaled with movement speed, most prominently under visual contact in the different-music condition (by contrast, the modulation in the same-music condition was less apparent, likely obscured by concurrent neural processes not considered in the ERP analysis, such as those related to coordination). The increased ERP amplitudes for faster compared to slower movements (Fig. 5) further highlight a well-established physiological property of sensory ERPs: their sensitivity to variations in stimulus intensity (Somervail et al., 2021).
***(IV) Social coordination.*** ERPs time-locked to changes in social coordination were associated with quadriphasic EEG modulations in occipital regions across all conditions. However, ERP amplitude differences between changes to in-phase *vs* anti-phase coordination emerged only when participants could see their partners and listened to same-tempo music (Fig. 5). The response pattern appears to bridge motor control and movement observation processes, showing a triphasic N-P-N sequence, similar to self-generated movement ERPs, followed by a positive occipital peak at 160 ms post-onset—resembling the inverted polarity of the posterior N160 observed for other-generated movements. The presence of a clear pattern in non-visual conditions suggests that these ERPs partially reflect motor activity, as changes in coordination coincide with movement initiation by either the self or the partner, a confound that traditional ERP analysis fails to fully resolve, unlike TRF analysis. Supporting this interpretation, ERP amplitude in non-visual conditions did not vary between changes to in-phase and anti-phase (Fig. 5), and the TRFs—designed to disentangle concurrent processes—did not reveal any EEG activity related to coordination, beyond motor activity, in the non-visual conditions (Fig. 3b).

**Figure 5.**
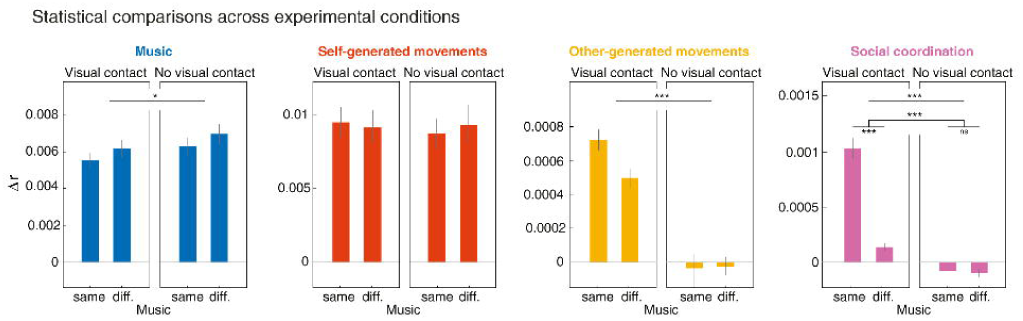
Event-related potential (ERP) analysis. ERPs evoked by salient changes in music, self-generated movements, other-generated movements, and social coordination. EEG time-series were epoched to peaks of spectral flux for music, peaks of velocity magnitude for self- and other-generated movements, and changes between in-phase and anti-phase for social coordination. ERPs amplitudes were compared across two groups of epochs within each variable: loud vs. soft sounds for music (Fz), fast vs. slow movements for self- (averaged across C3, C4) and other- (Oz) generated movements, and changes to in-phase vs. to anti-phase for social coordination (Oz). Grand-average ERPs are shown for the two groups of epochs within each variable and across all experimental conditions. Colored shaded areas represent ±1 SEM, while grey shaded regions highlight significant differences in ERP amplitude between groups of epochs at a given time point (permutation test over time, at the electrode of interest, cluster-corrected). Topographical maps display amplitude differences across electrodes within the time windows of identified clusters.

**Figure 6.**
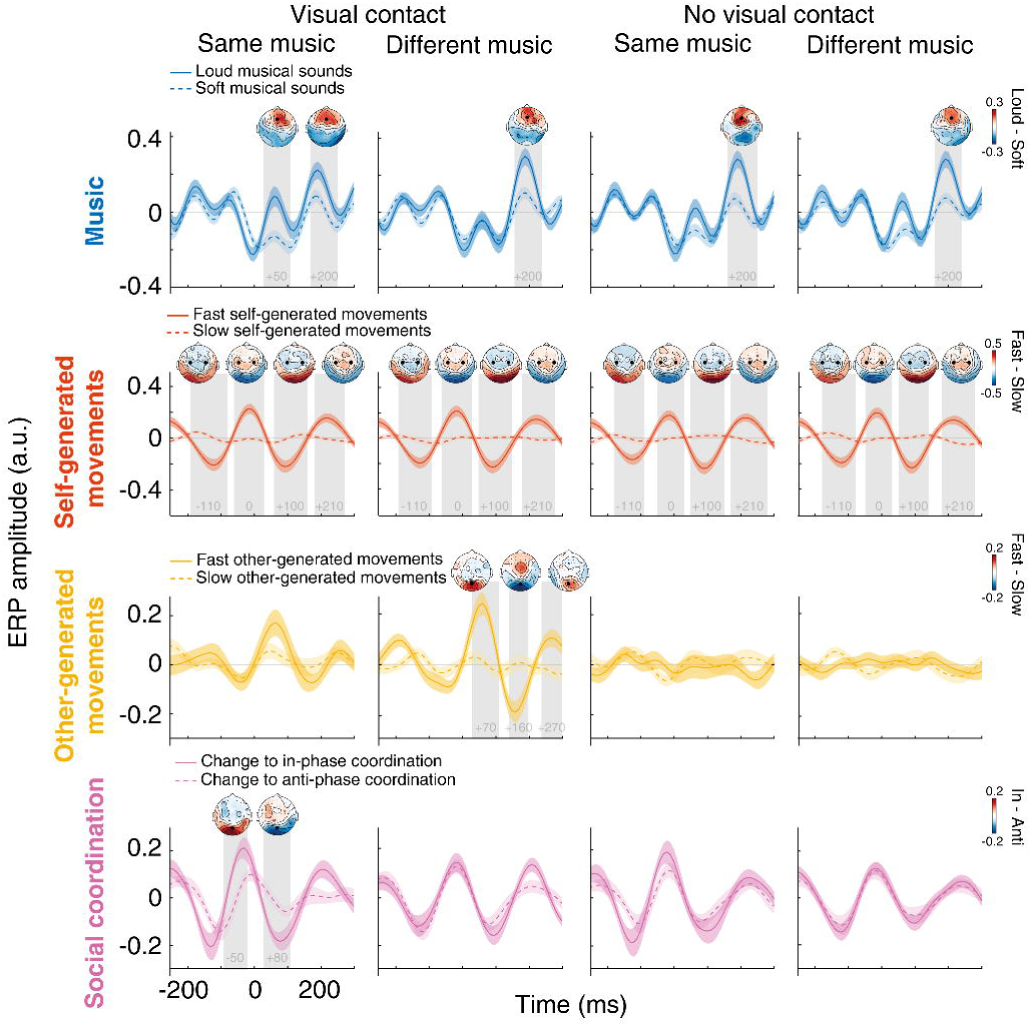
mTRFs tease apart body-part-specific motor activity. **a**, Topographical maps of the unique contribution of (self-generated) left- and right-hand movements to the predicted EEG. Δr topographical maps represent the grand-average difference in EEG prediction accuracy between the reduced models (excluding the body part of interest) and the full model (including all body parts, plus the neck control variable), for each EEG electrode and across all trials, regardless of experimental condition. We ran the TRF models without considering experimental conditions, given that no statistical difference was found across conditions in our main analysis (see Fig. 4). Separate TRF models for hands were derived by excluding each marker (‘L Hand’ or ‘R Hand’) from the full model. **b,** Same as (a), but for left- and right-foot movements. Separate TRF models for feet were derived by excluding each marker (‘L Foot’ or ‘R Foot’) from the full model. **c,** Same as (a) and (b), but for head movements. The head TRF model was derived by excluding all four head markers together.

### mTRFs tease apart body-part-specific motor activity

Thus far, the EEG activity related to self-generated movements was extracted using the velocity time-series of bounce movements (see Methods), either to predict EEG signal (mTRF analysis) or to time-lock EEG epochs (ERP analysis). To determine whether specific body parts contribute to distinct motor activities, we performed an additional TRF analysis using velocity time series associated with five distinct body parts: left and right hands, left and right feet, and head. Rather than relying on principal movements extracted from PCA, we modeled EEG signals using the kinematics of these specific body parts as input variables in the TRF models (Fig. 6). The unique contributions of the left and right hands (beyond that of all other body parts) to the EEG prediction exhibited lateralized spatial maps at central sites (Fig. 6a), a typical marker of hands’ motor control (Kornhuber and Deecke, 1965; Deecke et al., 1969; Shibasaki et al., 1980; Gerloff et al., 1997; Smulders and Miller, 2011; Vercillo et al., 2018; O’Neill et al., 2024). Furthermore, feet movements exhibited a more posterior topographical activation than hands (Fig. 6b), reminiscent of EEG differences found when comparing motor activity across hands and feet (Brunia et al., 2011). Notably, these feet-related EEG activities showed no clear lateralization, which is expected given the organization of the motor cortex (Gordon et al., 2023; O’Neill et al., 2024) and the limited spatial resolution of EEG (Osman et al., 2005). Indeed, as the feet are represented in the deeper, more central regions of the motor cortex, along the inner surface of the longitudinal fissure, it is notoriously difficult to differentiate EEG activity evoked by left vs. right foot movements (Osman et al., 2005; Jensen et al., 2023). Finally, head movements were associated with EEG activity not only in motor sites, such as C3 and C4 electrodes but also in occipital regions (Fig. 6c). Importantly, this occipital activation did not result from neck muscle artifact leakage, as neck

EMG’s contribution was already accounted for in the full model (see Methods). Moreover, no such occipital activation was found for hand or foot movements, suggesting that head movements specifically involve visual (besides motor) processing. Visual processes could be at play when moving the head (e.g., bouncing or head bobbing) as this involves salient changes in the field of view. Taken together, these findings support the conclusion that our TRF and ERP analyses (see Figs. 3 and 5) efficiently isolated neural processes related to self-generated movements. Moreover, beyond validating these prior results, this new analysis demonstrates the feasibility of isolating motor activity of specific body parts (note that in the prior analyses, motor activity related to bounce [involving all body parts] was assessed, limiting visibility into body-part-specific motor activity).

### Social coordination encoding acts beyond self and other

#### Coordination beyond self and other

In previous analyses, we demonstrated that adding the social coordination variable to models including music, self-, and other-generated movements yielded a gain in EEG prediction, suggesting neural encoding of coordination (Figs. 3 and 4). To ensure this gain was not solely attributed to the inclusion of spatial direction features – i.e., up versus down phases of bounce movement inherent in the social coordination variable but absent in the self- and other-generated movement variables – we conducted a supplementary mTRF analysis that included the spatial directions of both self- and other-generated movements. This analysis yielded cross-condition differences in unique contributions (i.e., Δ*r*) that were identical to those observed in our primary analysis (compare Fig. 7a, top, with Figs. 3a and 4), along with consistent model weights (compare Fig. 7a, bottom, with Fig. 3b). These findings indicate that the encoding of social coordination extends beyond merely representing the spatial directions of self- and other-generated movements in isolation. These results further suggest that the encoding of coordination is not merely driven by a modulation of the partner-evoked visual processes as a function of whether the self is moving congruently or not congruently with the partner (hence suppressing or amplifying the observed movements relative to the field of view). To further show that the encoding of coordination was not solely capturing this, we conducted an additional control analysis for which we re-referenced the other-generated movements to the position of the self. Even following such re-referencing, social coordination yielded a significant and unique contribution to EEG recorded from occipital sites, and this contribution was most pronounced under conditions of visual contact and shared music. Taken together, these findings indicate that the reported encoding of coordination is linked to a high-order process tracking the alignment between self- and partner-related movements, independently of the encoding of self and other taken alone.

**Figure 7.**
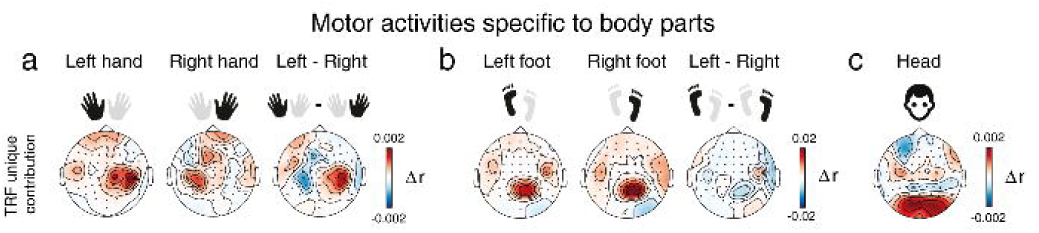
Tracking of social coordination beyond self and other. **a**, Results of the mTRF models associated with social coordination, incorporating spatial directions of self- and other-generated movements. Top: Social coordination uniquely predicted EEG activity at similar electrode sites than in the main analysis (Fig. 3a). Statistics revealed the exact same differences in unique contribution as observed in our main analysis (Fig. 4). Bottom: TRF regression weights exhibited similar patterns as in the main analysis (Fig. 3b). **b,** Coordination-related ERPs time-locked to self-generated (top) or other-generated (bottom) movement changes, at electrode Oz. ERPs related to changes to in-phase and anti-phase coordination are represented by continuous and dashed lines, respectively. Grand-average ERPs are shown for the two groups of trials associated with each variable across all experimental conditions. Colored shaded areas represent ±1 SEM, while grey shaded regions highlight significant differences in ERP amplitude between groups of trials at a given time point (permutation test over time, at Oz, cluster-corrected). Topographical maps display amplitude differences across electrodes within the time windows of identified clusters.

#### Coordination ERPs are time-locked to other-generated movements

“Coordination ERPs” were extracted by epoching EEG time-series to shifts from anti-phase to in-phase coordination and, vice versa, from in-phase to anti-phase. Here we investigated how such ERPs changed as a function of whether the shifts were driven by changes in movement direction produced by either the self (movement production) or the other (movement observation) (see Methods). Our analysis indicated that the EEG modulations previously associated with changes to in-phase coordination, specifically observed at occipital sites (Oz) and specifically under conditions of visual contact and same-tempo music (Fig. 5), were present only when these changes were time-locked to other-generated movement changes (Fig. 7b). This indicates that larger amplitude ERPs are evoked when a partner initiates a change in movement direction that leads to in-phase coordination compared to a change in movement direction that leads to anti-phase coordination. This result further strengthens the conclusion that the brain encodes social coordination and that this encoding is specifically driven by movements of the partner being in phase vs. anti-phase concerning self-initiated movements.

## Discussion

This study demonstrates that the neural processes underlying dance—a complex, natural, and social behavior—can be effectively isolated from EEG signals recorded from dyads dancing together. Using multivariate TRF models applied to dual EEG and full-body kinematics, we disentangled intertwined neural processes, separated them from movement artifacts, and confirmed their physiological origins through ERP analyses. This approach delineated sensory and motor processes underlying free-form, naturalistic dance: (I) auditory tracking of music, (II) control of self-generated movements, and (III) visual monitoring of partner-generated movements. Crucially, we also uncovered a previously unknown neural marker of social processing: (IV) visual encoding of social coordination, which emerges only when partners can make visual contact, is topographically distributed over the occipital areas, and is driven by movement observation rather than initiation. Additionally, movement-specific models highlighted “bounce” as the primary dance move driving EEG activity associated with both self-generated movements and movements observed in the partner. Together, these findings illustrate how advanced neural analysis techniques can illuminate the mechanisms supporting complex natural behaviors.

## mTRFs can unravel the complex orchestration of natural behavior

Recent advancements in mobile imaging and de-noising techniques have enhanced our ability to study neural activity during real-world behavior (Bateson et al., 2017; Niso et al., 2023). However, disentangling the contribution of the multiple simultaneous neural processes remains challenging. In this study, we addressed this challenge within the context of a spontaneous, interactive, yet controlled task, balancing ecological validity with experimental control (D’Ausilio et al., 2015). Using mTRFs, we successfully isolated four distinct, yet overlapping, neural processes underlying dyadic dance. ERP analyses confirmed that mTRF modeled responses, or model weights, align with well-characterized EEG potentials linked to sensory perception and motor control. This result suggests that mTRFs can capture physiologically established signals, akin to ERP analyses, but in real-world scenarios with multiple concurrent activities—contexts where traditional ERP approaches fall short.

ERP analyses struggle to isolate the unique contributions of individual processes amidst overlapping neural activities. This limitation is evident in our results: visual ERP modulation to movement speed was weak under visual contact and same-music conditions (Fig. 5), while mTRFs captured robust visual tracking (Figs. 3 and 4). This discrepancy likely stems from social coordination activity, which ERP analysis cannot disentangle, and which was particularly prominent in these conditions. Similarly, coordination ERPs appeared in non-visual conditions (Fig. 5), whereas mTRFs showed no corresponding activity, likely reflecting unaccounted self-motor contributions in ERP analyses. Although techniques like frequency tagging have addressed some of these challenges (Varlet et al., 2020, 2023; Cracco et al., 2022), they are limited to identifying periodic EEG responses and typically focus on univariate kinematics (e.g., gait cycles or hand trajectories). In contrast, mTRFs offer a precise characterization of neural responses to diverse features, effectively separating them from concurrent activities.

## The interplay between music and partner tracking

Dyadic dance requires simultaneous sensory processing of music and a partner’s movements, both of which contribute to coordinated behavior (Bigand et al., 2024). To what extent do these concurrent streams of information influence EEG activity, and how are these effects modulated by social factors like visual contact? Our findings show that model weights and ERPs associated with music, partner movements, and coordination exhibit similar amplitude, suggesting that each element—whether a musical sound, observed movement, or change in coordination—elicits an EEG response of comparable magnitude. Notably, visual processes accounted for less variance at occipital sites than auditory processes at frontal sites (see Δ*r* scales in Fig. 3a). This may reflect the broader range of EEG signals in occipital regions, which likely include visual processing of not only partner movements but also other visual cues and, importantly, artifactual leakage from neck movements (see Fig. 3a).

In visual-contact conditions, where both music (acoustic) and partner (visual) information were present, we observed a decrease in music tracking (Fig. 4). This reduction may arise from competition between visual and auditory modalities for attentional resources (Woods et al., 1992; Lavie, 2005; Molloy et al., 2015), especially in naturalistic dance, where both auditory and visual inputs drive coordination (Bigand et al., 2024). Naturalistic dance likely places heightened demands on visual input, as recent findings suggest that visual drivers dominate full-body rhythmic synchronization—a phenomenon not observed in simpler tasks like finger tapping (Nguyen et al., 2024).

## Movement control and observation

Our principal component analysis of dance kinematics revealed that bounce movements accounted for most EEG activity associated with self-generated movements (Fig. 2). Intriguingly, these movements predicted EEG activity not only over motor areas (e.g., electrodes C3 and C4), but also at occipital sites (Fig. 3a). To better understand these activities, we further dissected the components of bounce control, pinpointing motor activity specific to different body parts (Fig. 6). In participants engaged in free-form dancing, we successfully replicated established EEG findings observed during isolated movements, with more posterior-medial activity associated with foot movements and more central-lateralized activity for hand movements (Brunia et al., 2011). Notably, our analysis showed that head displacement was linked to occipital brain activity in addition to central motor activity, likely due to visual responses resulting from changes in the visual field (Testard et al., 2024). This analysis clarifies why the main mTRF for self-generated bounce movements included activity at occipital sites (Fig. 3a), suggesting that bouncing not only involves motor activity but also induces significant visual changes.

Bounce also emerged as the movement most predictive of EEG activity linked to visual tracking of a partner’s movements. This finding raises an intriguing question: what makes bounce particularly captivating compared to other dance movements? Our previous research has highlighted bounce’s key role in fostering interpersonal coordination (Bigand et al., 2024), serving as a supramodal (audio-visual) pace-setter between participants and their partners. This may explain why bounce is so prominent in predicting EEG activity associated with movement observation. This finding also suggests that EEG signals are particularly sensitive to salient movement changes, rather than merely high-amplitude movements. Indeed, while bounce explained less than 1% of the total kinematic variance (ranking 10th in the PCA), it accounted for over 80% of the EEG variance. This likely reflects bounce’s heightened salience, possibly driven by the fact that this movement was the only one peaking sharply with each musical beat (Bigand et al., 2024).

## Encoding of social coordination

Our study reveals that coordination between self- and other-generated movements uniquely predicts EEG signals recorded at occipital electrodes. Recent research in social neuroscience has shown that EEG can delineate separate components supporting coordinated behaviors: some monitor self- and partner-generated actions distinctly, while others integrate the joint action outcome produced by oneself and the partner (Novembre et al., 2016; Varlet et al., 2020). In line with this, we identified three distinct neural processes—control of one’s own movements, observation of a partner’s movements, and processing of social coordination (Fig. 3)—and observed heightened coordination tracking in conditions where musical synchrony between participants was greater. Importantly, our findings suggest that the encoding of coordination does not merely combine the individual “self” and “other” processes; rather, it captures a distinct neural representation of their coordination (see Results and Fig. 7a).

The temporal response underlying social coordination tracking integrates both motor control (self) and movement observation (other), as evidenced by the N-P-N pattern and a subsequent modulation peaking around 160 ms. Notably, this response appears to be triggered by observing a partner’s movements, rather than initiating one’s own actions. ERPs associated with changes in the partner’s movements—rather than self-initiated actions—were modulated by social coordination at visual sites (see Results and Fig. 7b). This finding suggests that neural tracking of coordination is more reliant on visual monitoring of the partner than on the internal control of one’s own movements, aligning with earlier observations that this process is localized in visual areas and enhanced during visual contact.

In summary, we identified a previously unknown neural marker of social processing, with five key observations: 1) it is topographically distributed over the occipital areas; 2) it emerges when participants can see each other and is most pronounced when musical synchrony between them is high; 3) its underlying neural signal integrates components from both self and other processes; yet 4) rather than merely combining the individual “self” and “other” components, it represents a distinct neural encoding of their coordination; and 5) it is primarily anchored to movement observation, not movement initiation.

## Bridging traditional physiology with real-world applications

Our findings show that neurophysiological signals, traditionally examined in controlled settings, can be disentangled and analyzed within real-world contexts. This highlights the potential for future research to incorporate ecologically valid stimuli and behavioral predictors (e.g., body movements, eye gaze, speech) into multivariate modeling. Such an approach could deepen our understanding of brain processes during live social interactions—a field of growing significance across human adult (Dumas et al., 2010; Pan et al., 2018; Koul et al., 2023; Cross et al., 2024; Orgs et al., 2024), developmental (Wass et al., 2018, 2020; Nguyen et al., 2020, 2021, 2023) and animal studies (Zhang and Yartsev, 2019; Rose et al., 2021; Yang et al., 2021).

## Acknowledgments

F.B., S.F.A., and G.N. are supported by the European Research Council (ERC, MUSICOM, 948186). R.B. is supported by the European Union (MSCA, PHYLOMUSIC, 101064334). T.N. is supported by the European Union (MSCA, SYNCON, 101105726). We thank Alison Rigby for her help with data collection, and Raoul Tchoï for his help creating the stimuli.

## Code accessibility

All original code is publicly available on Github repositories: https://github.com/felixbgd/dancing_brain

## Data accessibility

EEG, EMG, EOG, kinematic, and musical data have been deposited to IIT Dataverse and are publicly available as of the date of publication: https://dataverse.iit.it/privateurl.xhtml?token=bb0689c8-137d-4742-9f94-d7c6b0148827

## Multimedia

**Video 1. The principal (dance) movements, related to Fig. 2**

Video showing original movement data (left) and their decomposition into 15 principal movements (PMs) explaining >95% of the kinematics variance (right). Representative data are displayed (excerpt from a single trial, corresponding to when participants listened to the full refrain of the song). For the sake of clarity, the PMs are animated with different levels of exaggeration (i.e. the PM scores were amplified by a factor of 1.5 (PM3), 2 (PM4,7,9,11,15), 3 (PM8), or not amplified (all other PMs)). The PMs are reminiscent of common dance moves (spelled out in italics).

